# The E3 ubiquitin ligase UBR5 regulates centriolar satellite stability and primary cilia formation via ubiquitylation of CSPP-L

**DOI:** 10.1101/090100

**Authors:** Robert F. Shearer, Kari-Anne Myrum Frikstad, Jessie McKenna, Rachael A. McCloy, Niantao Deng, Andrew Burgess, Trond Stokke, Sebastian Patzke, Darren N. Saunders

## Abstract

Primary cilia are crucial for signal transduction in a variety of pathways, including Hedgehog and Wnt. Disruption of primary cilia formation (ciliogenesis) is linked to numerous developmental disorders (known as ciliopathies) and diseases, including cancer. The Ubiquitin-Proteasome System (UPS) component UBR5 was previously identified as a putative modulator of ciliogenesis in a functional genomics screen. UBR5 is an E3 Ubiquitin ligase that is frequently deregulated in tumours, but its biological role in cancer is largely uncharacterised, partly due to a lack of understanding of interacting proteins and pathways. We validated the effect of UBR5 depletion on primary cilia formation using a robust model of ciliogenesis, and identified CSPP1, a centrosomal and ciliary protein required for cilia formation, as a UBR5-interacting protein. We show that UBR5 ubiquitylates CSPP1, and that UBR5 is required for cytoplasmic organization of CSPP1-comprising centriolar satellites in centrosomal periphery. Hence, we have established a key role for UBR5 in ciliogenesis that may have important implications in understanding cancer pathophysiology.

## Introduction

Primary cilia form at the surface of most mammalian cell types, and have been implicated in sensory perception, cell signalling and development. Primary cilia comprise a microtubule axoneme and ciliary membrane extending from the basal body (Satir and Christensen, 2007), which is surrounded by granular structures known as centriolar satellites (Lopes et al., 2011) largely composed of percentriolar material 1 (PCM1) and other key proteins (Kubo et al., 1999). In contrast to motile cilia, primary cilia lack the core microtubule/dynein structure that provides motility to the former (Roth et al., 1988). During mammalian development, primary cilia are present from around embryonic day E6.0, and are retained throughout gestation in a lineage dependant manner (Bangs et al., 2015). Multiple signalling pathways including Hedgehog (Hh) require intact cilia for effective signal transduction (May et al., 2005), and as such understanding the dynamics of ciliogenesis and cilia function is highly relevant to understanding development and cancer (Michaud and Yoder, 2006). Recent characterisation of the complex ciliary proteome and interactome (Mick et al., 2015, Pazour et al., 2005, Boldt et al., 2016, Wheway et al., 2015, Gupta et al., 2015), demonstrate the highly organised and dynamic structure of primary cilia.

Ubiquitylation is one of the most common protein post-translational modifications, resulting in attachment of one or more Ubiquitin (Ub) molecules to substrate proteins via an ATP-dependent enzymatic cascade known as the Ubiquitin-Proteasome System (UPS) (Ciehanover et al., 1978, Hershko et al., 1981). The various signalling outcomes of protein ubiquitylation are determined by the topology of the attached poly-Ub chains (Passmore and Barford, 2004). For example, K48 linked poly-Ub targets substrates for proteasomal degradation, but alternate linkages (K63, K29 etc) are emerging as key regulators of cell signalling via modulation of protein function, localisation and protein-protein interactions. Key proteins associated with tumour initiation (e.g. p53, NF-ĸB) are regulated by the proteasome, and proteasome inhibitors such as PS-341 and Bortezomib (Velcade) are under trial or approved for clinical use in hematological malignancies (Johnson, 2014, Orlowski et al., 2002).

Integrity of centriolar satellites are crucial for centrosome related functions, including ciliogenesis (Tollenaere et al., 2015). The UPS is emerging as a key regulator of ciliogenesis and centriolar satellite stability (Shearer and Saunders, 2016). E3 Ub ligases, which largely determine substrate specificity and are an important rate-limiting component of the UPS, can regulate expression of various proteins crucial to ciliary axoneme extension. For example, the Cullin Ub ligase MIB1 regulates multiple centriolar proteins, including PCM1 (Villumsen et al., 2013, Cajanek et al., 2015) and extension of the axoneme is initiated by UPS-mediated degradation of the ciliogenesis inhibitor Trichoplein (Inoko et al., 2012, Kasahara et al., 2014). A genome-wide RNA interference (RNAi) screen for regulators of ciliogenesis identified a number of components of the UPS (Kim et al., 2010), including the E3 Ub ligase UBR5 (ubiquitin protein ligase E3 component N-recognin 5). UBR5 is a highly conserved gene (Callaghan et al., 1998, Mansfield et al., 1994) required for normal embryonic development, correct functioning of cell cycle checkpoints, and multiple aspects of the DNA damage response (Shearer et al., 2015). UBR5 is frequently de-regulated in many cancer types by amplification and/or mutation (Meissner et al., 2013, Clancy et al., 2003, O’Brien et al., 2008), but the full complement of UBR5 substrates, and hence the mechanistic role of UBR5 in cancer, remain to be defined.

We sought to further investigate the role of UBR5 in ciliogenesis in a cancer context. We show that UBR5 regulates primary ciliogenesis via a novel interaction with, and ubiquitin-mediated sub-cellular localisation of the centriolar satellite protein CSPP1. This identifies UBR5 as a novel ciliary regulator, with implications for understanding cell signalling in development and cancer.

## Materials and Methods

### Gene expression correlation

Global gene expression correlation in the NCI-60 panel of cancer cell lines (n=60, (Stinson et al., 1992)) was analysed using the Pattern comparison tool from CellMiner (Reinhold et al., 2012). Expression intensity Z-scores for 26065 genes were correlated against *UBR5* expression based on Affymetrix microarray transcript intensity level. Significance based Pearson’s correlation co-efficient with p<0.05 without multiple comparisons. Linear regression analysis was performed comparing intensity Z-scores for *UBR5* and *CSPP1* expression using data obtained from the Cross-correlations tool from CellMiner. Co-expression data was validated using the Cancer Genome Atlas (TCGA) (Ciriello et al., 2015), Cancer Cell Line Encyclopaedia (CCLE) (Barretina et al., 2012) and the Molecular Taxonomy of Breast Cancer Interational Corsortium (METABRIC) (Curtis et al., 2012) cohorts. Analyses were performed using cBIOPortal (Cerami et al., 2012, Gao et al., 2013). Co-expression measured on mRNA expression determined using RNA-seq data (Z-score, threshold +/−2.0).

### Plasmids

Full length *UBR5* ORF was obtained from pEGFP-C1 EDD (Addgene #37190) (Henderson et al., 2002) and used to create pENTR221-UBR5 (Addgene #81062). *UBR5* cDNA was digested from pEGFP-C1 *EDD* and was cloned into pENTR221 containing synthetic fragments (GeneArt, Life Technologies) of the 5’ and 3’region (bp 1-1129 and 6933-8404 respectively) of *UBR5*. Mutagenesis of *UBR5* was achieved by subcloning synthetic fragments (GeneArt, Life Technologies) of the HECT domain (bp 6933-8404) with a mutation (bp t8302g and g8303c) corresponding to C2768A (herein ΔHECT, Addgene, #81065), the UBR5 MLLE domain (Kozlov et al., 2007) (bp 6933-8404) with mutations (bp t7204g, a7205c, t7206c, a7243g, a7244c and a7245c) corresponding to Y2402A and K2415A (herein ΔMLLE, Addgene, #81064), and the UBR domain (bp 3163-4217) with a mutation (bp g3704t) corresponding to W1235L (herein ΔUBR, Addgene, #81063). A Gateway entry vector encoding full length *CSPP-L* was generated by PCR using modified flanking primers and the previously described pCSPP-L-EGFP vector (Patzke et al., 2006).

Gateway entry vectors were used to generate expression clones using the following destination vectors: Vivid Colors^™^ pcDNA^™^6.2/N-EmGFP-DEST (V356-20, Invitrogen), Bi-molecular fluorescence complementation (BiFC) vectors pDEST-V1-ORF (Addgene 73635) and pDEST-V2-ORF (Addgene 73636) (Croucher et al., 2016). GFP-UBR5 (Addgene 52050) and GFP-UBR5 ΔHECT (52051, Addgene) expression vectors are described (Gudjonsson et al., 2012). V1-Ub and V2-Ub expression vectors are described (Lee et al., 2015). Recombination was catalysed by Gateway^®^ LR clonase II enzyme mix (11791-020, Invitrogen) according to manufacturer’s instructions.

Short-hairpin RNA (herein shRNA) sequences to UBR5 were obtained from the RNAi codex project (Olson et al., 2006). Haripins were cloned into pEN_TmiRc3 (25748, Addgene), before shuttling into pSLIK Gateway^®^ compatible expression vectors encoding Venus (25734, Addgene) or G418 (25735, Addgene) selectable markers as described (Shin et al., 2006). The hairpin sequence 5’-CGCAGTGAATGTAGATTCCAAA-3’ (HP_6400, herein shUBR5 (Addgene, #81066) was found to efficiently deplete UBR5 and was used for experiments. A scrambled sequence 5’-TCGATGCTCTAAGGTTCTATC-3’ (herein shNT, Addgene, #81067) was used as a non-targetting control. pLV-CCN-H2B-mCherry was a kind gift from Marc Giry-Laterriere.

### PAGE and Immunoblot

All lysates were made using RIPA buffer supplemented with protease inhibitors (1183617001, Roche Diagnostics), and 10mM N-Ethylmaleimide (E3876-5G, Sigma). Cultured cells were washed with PBS and scraped from culture vessels in the presence of lysis buffer on ice. Lysates were cleared by centrifugation at 4°C and the total protein concentration determined using Protein Assay Dye Reagent (500-0006, Bio-rad) according to manufacturer’s instructions. Samples were separated using SDS-PAGE, transferred to Immobilon-P PVDF 0.45μm membrane (IPVH00010, Merck Millipore) and subsequently immunoblotted using standard procedure. Densitometry was performed using ImageJ. Lane density was plotted and relative band intensity determined by area under curve analysis. Densitometry was restricted to comparison of lanes from the same exposure and run on the same gel. Intensity was normalised to loading control and was standardised to the first lane of each gel. Theoretical size of fusion proteins was calculated using Compute pI/Mw tool available on the ExPASy server (Bjellqvist et al., 1993) via the average resolution setting.

The following antibodies were used for immunobloting. Goat anti-EDD N-19 (sc-9561, Santa Cruz) diluted 1:5000. Rabbit anti-EDD1 (A300-573A, Bethyl Laboratories) diluted 1:5000. Rabbit anti-CSPP1 (Binds CSPP-L only, 11931-1-AP, Proteintech) diluted 1:5000. Rabbit anti-PCM1 (ab72443, Abcam) diluted 1:5000. Mouse anti-GFP (MMS-118P, Covance) diluted 1:5000. Mouse anti-GFP (11814460001, Roche) diluted 1:5000. Rabbit anti-Ubiquitin (ab7780, Abcam) diluted 1:2000. Mouse anti-β-ACTIN (A5441, Sigma-Aldrich) diluted 1:50000. Mouse anti-GAPDH (ACR110PT, Acris-Antibodies) diluted 1:10000. Rabbit anti-Ubiquitin (ab7780, Abcam) diluted 1:2000. Rabbit anti-Ubiquitin (linkage specific K48) (ab140601, Abcam) diluted 1:5000. Rabbit anti-Ubiquitin (linkage-specific K63) (ab179434, Abcam) diluted 1:5000. Rabbit anti-γ-tubulin (T3320, Sigma-Aldrich) 1:5000. Mouse anti-γ-tubulin (T6557, Sigma-Aldrich) diluted 1:5000. Rabbit anti-CEP290 (ab85728, Abcam) diluted 1:2000. All antibodies were diluted in 5% (w/v) BSA TBS solution.

### Immunoprecipitation

GFP-tagged fusion proteins were isolated from WCE using GFP-Trap (GTA-100, Chromotek) affinity purification reagent according to manufacturer’s instructions. In brief, 20μl of bead slurry was washed twice in 10mM Tris-HCl (pH 7.5) with 150mM NaCl and 0.5mM EDTA before addition of 250μg WCE. Samples were incubated for 1 hour at room temperature with gentle end-over-end mixing. Beads were washed twice, and bound proteins eluted by heating at 95°C for 10 minutes in 1XSDS gel loading dye.

### Cell culture

The following cell lines were cultured at 37°C with 5% CO2 and passaged according to ATCC recommendations. Cell line identity was verified using standard in house authentication. hTERT-RPE1 (CRL-4000, ATCC) human hTERT immortalised retina pigmented epithelial cells were maintained in DMEM-F12 culture medium (31331-028, Life Technologies) supplemented with 10% FBS(v/v) and 1% Penicillin-Streptomycin solution (v/v) (P4333, SIGMA). Human embryonic kidney (HEK293T, CRL-3216, ATCC) cells were grown using DMEM culture medium (11995-065, Life Technologies) supplemented with 10% FBS (v/v), MEM Non-essential amino acids (11140-050, Life Technologies) and sodium pyruvate (11360-070, Life Technologies). MDA-MB-231 (HTB-26, ATCC) human mammary carcinoma cells were grown with RPMI 1640 culture medium (11875-085), Gibco) supplemented with 10% (v/v) FBS, 10mM HEPES buffer (15630-080, Life Technologies), 0.2 IU/ml Human Insulin. Inhibition of proteasomal degradation was achieved where indicated using 10μM MG-132 (474790-1MG, Calbiochem) in complete growth media for prior to harvest. HEK293T cells expressing H2B-mCherry were generated by stable transduction of pLV-CCN-H2B-mCherry viral supernatant. Cells expressing H2B-mCherry were grown under 500μg/ml G418 selection and sorted for moderate expression as described (McCloy et al., 2014). HEK293T cells expressing a short hairpin to UBR5 (shUBR5) or a non-targetting scrambled control (shNT) were generated by stable transduction of pSLIK (see above) viral supernatant. Viral transduction to achieve minimal MOI performed visually in serial dilution as described (Shearer and Saunders, 2015). Sorting performed by Garvan Flow Cytometry Facility staff.

Plasmid transfections were performed using Xtreme Gene 9 HP transfection reagent (06366236001, Roche). Cells were plated out 24 hours prior to achieve roughly 50% confluency at the time of transfection. Plasmid DNA was mixed with transfection reagent in a 1:3 (μg DNA:μl reagent) ratio, diluted in a total volume of 100μL Opti-Mem I reduced serum media (31985-070, Life Technologies). Complexes were incubated for 15 minutes at room temperature before being added to cells with complete growth media. Transient gene silencing in HEK293T cells using siRNA was performed using Lipofectamine 2000 (11668019, Life Technologies) according to manufacturer’s instructions. 5×10^5^ cells were plated in a 6-well plate 24 hours prior to transfection. 25pmol of siRNA was combined with 7μl transfection reagent diluted in 100μl total volume. After plating, cells were subject to imaging for proliferation assay (see below) before harvest and lysis at 24-48 hours as indicated. Epifluorescence imaging was performed using a Leica DM550 microscope.

### Ciliogenesis assay

5×10^4^ cells were seeded on high precision glass coverslips (0.17 ± 0.01mm; 1014/10, Hecht Assistent) in 30mm wells, allowed to adhere overnight and transfected with siRNA using RNAiMAX (Life Technologies) for transient gene silencing. 48h posttransfection cells were washed twice in serum-free DMEM-F12 and cilia formation induced by continued incubation in serum-free medium for 48h. For cilia detection cells were fixed in methanol (-20C) and stained subsequently with a glutamylated tubulin specific mouse monoclonal antibody (AG-20B-0020-C100, Adipogen AS) and a mouse specific Cy3 labelled secondary antibody (715-165-151, Jackson Immuno Research) for labelling of ciliary axonemes. At least 150 cells were scored by manual inspection on an AxioImagerZ.1 epifluorescence microscope equipped with a 40x/NA0.95 and a 63x/NA1.4 Plan-Apochromat lens and a HXP120 Metal-Halide Illuminator (Zeiss).

### Immunofluorescence microscopy

Cells were grown on heat-sterilized cover glasses (No.1014; Glaswarenfabrik Karl Hecht GmbH & Co KG, Sondheim/Rhön, DE), fixed for 15 min in 1% neutral buffered formaline solution at room-temperature prior to post-fixation in methanol (-20C). Cells were re-hydrated for IFM staining by three consecutive washes in Phosphate buffered saline (PBS) and blocked and permeabilized for 15 min in PBS-AT (PBS containing 5% w/vol. Bovine serum albumine (Sigma-Aldrich) and 0.1% vol/vol Triton-X-100 (Sigma-Aldrich)). Cells were stained with primary antibodies for 2hrs at r.t., washed thrice in PBS, and stained with secondary antibodies for 1 hr. All antibody incubations were performed in PBS-AT. Cells were washed in thrice PBS, counterstained for DNA (Hoechst 33258 in PBS, Sigma), washed briefly in distilled water, dried and mounted on object glasses using Prolong Gold (Life Technologies, Carlsbad, CA, US). Fluorescence images were acquired using appropriate optical filters on an multifluorescent bead calibrated AxioImager Z1 ApoTome microscope system (Carl Zeiss, Jena, DE) equipped with a 100× or a 63× lens (both PlanApo N.A.1.4) and an AxioCam MRm camera. To display the entire cell volume, images are presented as maximal projections of z-stacks using Axiovision 4.8.2 (Carl Zeiss).

Images for quantitative IFM imaging were acquired on a multifluorescence submicron beads calibrated CellObserver microscope system (Carl Zeiss) equipped with a 40×/1.3 PlanApo Phase 3 lens and an AxioCam MRm camera. Images were acquired with constant exposure times at 10 random positions per coverslip and in seven optical sections at 0.5μm distance, centered around focal planes for cilia. Central focal planes were identified by γ-tubulin labeling as centrosome reference using a contrast based autofocus routine (AxioVision 4.8.2). Image analysis was performed in Fiji/ImageJ (Schindelin et al., 2012). Sum projections of of individual channels were background corrected using a 5px rolling circle algorithm and segmented by signal intensity and morphological thresholds. Thresholded γ-tubulin signals defined the centrosome compartment mask. The radius of the centrosome area was iteratively dilated (20x) to cover the pericentrosomal area, and subtracted for the core centrosomal area to create the centriolar satellite mask. Fluorescence signal intensities in thresholded areas under each mask were measured in all channels to obtain integrated signal intensities.

The following antibodies were used for immunofluorescence. Rabbit anti-PCM1 (ab72443 Abcam) diluted 1:1000. Mouse anti-γ-tubulin (T6557, Sigma) diluted 1:500. Rabbit anti-CSPP1 (Binds CSPP-L only, 11931-1-AP, Proteintech) diluted 1:500. Mouse anti-Glutamylated tubulin (GT335, Enzo Life Sciences) diluted 1:500. All statistical analyses were performed using Prism (Graphpad Software).

### Bi-Molecular Fluorescence Complementation (BiFC)

Bi-molecular Fluorescence Complementation (BiFC) allows fluorescent visualisation of binary protein-protein interactions (Kerppola, 2008). Proteins of interest were expressed fused to (either Met^1^-Gln^157^ (V1) or Lys^158^-Lys^238^ (V2)) separated by a 2xGGGGS linker sequence (Croucher et al., 2016). Recombination of the Venus fluorescent protein indicates a positive interaction. 5×10^5^ HEK293T cells (containing stable H2B-mCherry) were plated in a 6-well plate containing a round coverslip 24 hours prior to transfection. Cells were transfected with 500ng of each plasmid construct (see above) and incubated for 18 hours before coverslips were mounted with Vectashield mounting medium (H-1400, Vector Laboratories). Remaining cells were harvested for immunoblot analysis to ensure correct fusion protein translation. Confocal imaging was performed using a Leica TCS SP8 confocal microscope optimised for at least 75nm resolution. Gain and resolution were maintained across all samples within experiments. Fluorescent images were pseudocoloured with appropriate LUT and merged using ImageJ.

## Results

### Depletion of UBR5 confers concurrent disruption of centriolar satellites and inhibition of primary cilia formation

UBR5 was identified as a putative regulator of primary cilia formation in a functional genomics screen (Kim et al., 2010) along with a number of other UPS components (Shearer and Saunders, 2016). We validated the effect of UBR5 depletion on cilia formation using a robust ciliogenesis assay (Fig 1A) and examined the functional significance of UBR5 depletion in the context of cilia/centriolar satellite stability/organization (Fig 1B). siRNA-mediated depletion of UBR5 almost completely attenuated primary cilia formation in hTERT-RPE1 cells (i.e. observed in fewer than 20% of cells), as determined by glutamylated tubulin stain (Fig 1C,D). Furthermore, centriolar satellites were dispersed with UBR5 depletion (Fig 1C,D). Levels of disruption were similar to decreases observed with depletion of PCM1 (Fig 1C,D), which recruits another E3 ligase MIB1 to the centrosome (Wang et al., 2016). Cilia were absent in 89% of hTERT-RPE1 with UBR5 depletion under serum starvation, and 94% of these cells displayed dispersed centriolar satellites (Fig. 1D). Approximately 77% of control hTERT-RPE1 cells expressed a primary cilium, of which 85% had distinct centriolar satellite formation around the basal body (Fig 1D).

**Figure 1:**
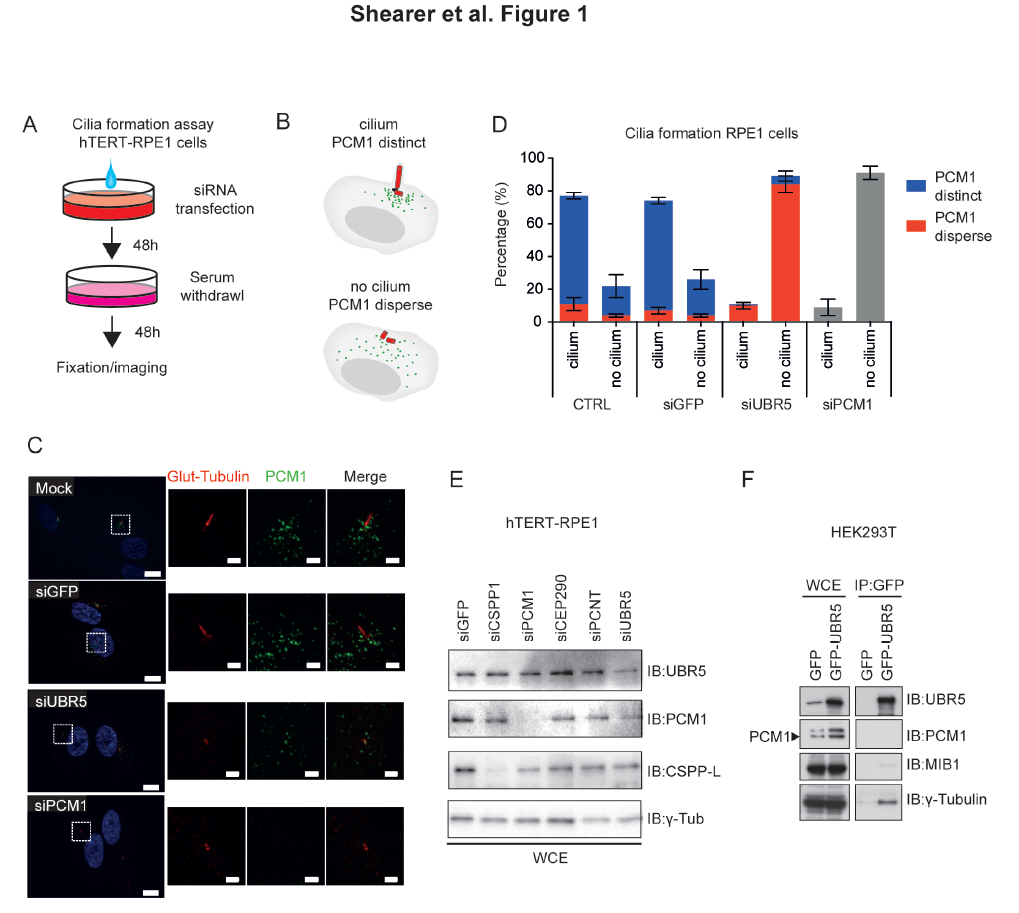
UBR5 depletion disrupts centriolar satellite stability and primary cilia formation in hTERT-RPE1 cells. **A**. Schematic describing ciliogenesis assay. **B**. Scoring criteria for ciliogenesis/centriolar stability **C**. Depletion of UBR5 attenuates primary cilia formation in and causes dispersion of centriolar satellites in RPE1 cells. Cilium and basal body stained with Glutamylated-tubulin (red), and satellites stained using PCM1 (green). Similar phenotypes are observed for depletion of PCM1. Nuclear marker Hoechst33258 (blue). High powered inset fields indicated by dashed box. Transfections performed ~72 hours prior to imaging. Data representative of 2 independent experiments, with 150 cells counted per condition per experiment. Confocal imaging, bar = 1μm in close-ups, 10μm in low power magnifications **D**. Quantitation of data depicted in (C) shows the strong penetrance of siUBR5 and siPCM1 phenotype. Cartoon depicts how staining was scored. **E**. Immunoblot showing siRNA efficacy in RPE1 cells used for staining in (C). **F** Co-immunoprecipitation of UBR5 and western blotting for potential interactions with PCM1, MIB1 and γ-tubulin (a marker of the centrosome). Transfections performed 48 hours prior to imaging.

Given this effect on UBR5 depletion on centriolar satellite stability, we hypothesised that UBR5 may act to stabilise centriolar material and investigated potential protein-protein interactions with known structural components of the centrosome and regulators of centriolar satellite stability. We performed pulldowns of GFP-tagged UBR5 in HEK293T cells, and observed co-immunoprecipitation of UBR5 and γ-tubulin, indicating that UBR5 can interact with structural components of the centrosome (Fig 1F). The UPS is known to regulate via ubiquitin-mediated regulation of PCM1 function by the E3 ubiquitin ligase MIB1 (Villumsen et al., 2013, Wang et al., 2016). We could not detect an interaction between UBR5 and PCM1, and we observed only a very weak putative interaction with MIB1 (Fig 1F). Furthermore, UBR5 depletion in HEK293T cells did not significantly alter levels of PCM1, MIB1 or γ-tubulin proteins (Fig. S1). Together, these data suggest that the role of UBR5 in ciliogenesis is not mediated via directly affecting PCM1/MIB1.

### Co-expression of CSPP1 and UBR5 in human cancer cell lines and primary cancer biopsies

To identify potential mechanisms by which UBR5 regulates cilia/centrosome biology, we analysed publicly available gene expression datasets to identify candidate genes co-regulated with *UBR5*. Coordinate expression of genes functioning in common pathways (i.e. synexpression groups) is a widespread phenomenon in eukaryotes and co-expression analysis has proven to be a powerful approach to identify novel gene function (Eisen et al., 1998, Wu et al., 2002, van Noort et al., 2003, Niehrs and Pollet, 1999). As *UBR5* is frequently altered in multiple cancer types (Shearer et al., 2015), expression data from the NCI-60 panel of cancer cell lines was utilised (Stinson et al., 1992). Analysis of mRNA co-expression in the NCI-60 panel using CellMiner (Reinhold et al., 2012) revealed a strong positive correlation between *UBR5* and a number of genes (Fig 2A) (Pearson correlation coefficient with n=60, significance cut-off at 0.254). Many genes co-expressed with *UBR5* are in close genomic proximity on chromosome 8q22 - a region of known genomic instability of solid tumours - and share common enhancer/repression elements (http://atlasgeneticsoncology.org/Indexbychrom/idxa_8.html). These genes were therefore excluded from further consideration.

**Figure 2:**
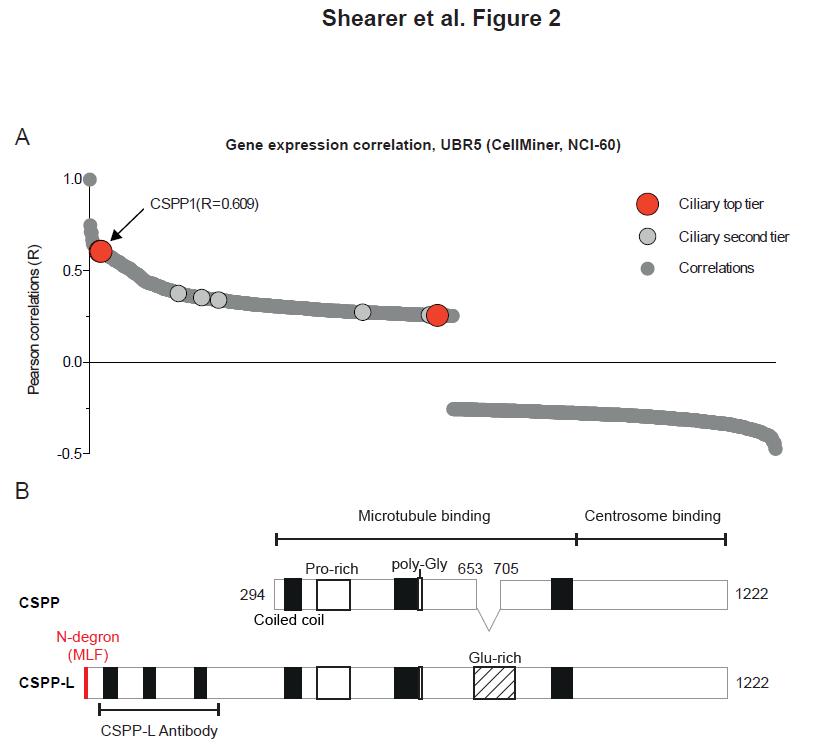
Co-Expression of UBR5 gene. **A**. Waterfall plot showing global correlation of gene expression against *UBR5* expression in NCI-60 cancer cell line panel, obtained via CellMiner (Reinhold et al., 2012). Cut-off set at Pearson correlation of 0.254. Putative ciliary proteins are indicated, with top tier proteins labelled and indicated with larger red dots, second tier proteins also labelled and shown with larger grey dots. **B**. Structure of CSPP1 protein isoforms. Note that the extra 294 amino acid N-terminal region of CSPP-L is the binding region of the selected CSPP1 antibody. N-degron motif shown in red. Coiled coil domains shown with a black box. Proline-rich domains shown with a white box. Glutamate-rich domains shown with a white striped box.

Of 855 genes significantly co-expressed with *UBR5* (Fig 2A), several have been previously identified as candidate ciliary localising proteins (Mick et al., 2015), including *Centrosome and Spindle Pole associated Protein 1 (CSPP1), YTH N(6)-methyladenosine RNA binding protein 3 (YTHDF3)* - which contains an RNA binding YTH motif (Stoilov et al., 2002), and *Septin 7 (SEPT7)* - which contains a GTP-binding motif (Serrao et al., 2011). The top-ranked candidate gene significantly co-expressed with *UBR5* in cancer cell lines was *CSPP1* (p<0.0001). CSPP1 is enriched in primary cilia (Mick et al., 2015, Patzke et al., 2010) and has established functional roles in regulation of spheroid formation, cell division and ciliogenesis (Asiedu et al., 2009, Patzke et al., 2010, Zhu et al., 2015, Sternemalm et al., 2015). Mutations in *CSPP1* have been implicated in ciliopathies (Tuz et al., 2014, Akizu et al., 2014, Shaheen et al., 2014) and expression of CSPP1 isoforms display distinct restriction in breast cancer subtypes (Sternemalm et al., 2014). The large and predominantly expressed isoform of CSPP1, CSPP-L, localises to the centrosome, where it is required for primary cilium formation in non-cycling cells (Mick et al., 2015, Patzke et al., 2010, Gupta et al., 2015), and dynamically re-localizes to the spindle apparatus of dividing cells, where it aids chromosome movements and cytokinesis during cell division (Patzke et al., 2006, Asiedu et al., 2009, Zhu et al., 2015).

Co-expression of *CSPP1* with *UBR5* in the NCI-60 cohort was confirmed by linear-regression analysis of NCI-60 cell line expression data (R^2^=0.3709, p<0.0001, Fig. S2A), and validated in independent breast cancer gene expression datasets from the TCGA (Fig. S2B, Table S1), CCLE, and METABRIC cohorts (Table S2). Co-expression of *CSPP1* with *UBR5* was evident in all cohorts, irrespective of subtype (Tables S1, S2). Over-expression of *UBR5* and *CSPP1* mRNA was the predominant form of altered expression, and co-amplification did not correlate with increased overall genomic instability (Fig S2B).

There are three known protein products of the *CSPP1* gene (Fig 2B). CSPP is a 101.5kDa protein that contains both microtubule and centrosome binding domains (Patzke et al., 2005). CSPP-L, contains an extra 294 amino acids at the N-terminus, and an additional 51 amino acids in the mid-domain, which affects how CSPP1 interacts with microtubules (Patzke et al., 2006). Of relevance to the ubiquitin context, the extended N-terminal region of CSPP-L harbours an N-degron motif (not present on CSPP) that likely targets CSPP-L for ubiquitylation by UBR-box E3 ubiquitin ligases (Tasaki et al., 2009). A third protein isoform of *CSPP1* comprising the C-terminal 379aa of CSPP-L is highly expressed in the nucleus of luminal breast cancer cells (Sternemalm et al., 2014). Western blot analysis of a panel of breast cancer cell lines showed co-expression of UBR5 and CSPP-L at the protein level (R^2^=0.2532, p=0.017, Fig. S2C,D), irrespective of subtype (Fig. S2E). Note that the epitope recognised by the CSPP1 antibody used in Fig. S2D is specific to the CSPP-L isoform.

### UBR5 regulates ubiquitylation of CSPP-L

In the context of UBR5 and CSPP-L co-expression, common functional roles in ciliogenesis, and a recently established interaction between PCM1 and CSPP-L, and localization to centriolar satellites (Sternelalm et al., under review), we hypothesised that UBR5 may regulate ciliogenesis via maintenance of centriolar satellite stability/organization and possibly involving CSPP-L. We therefore investigated a possible direct interaction between UBR5 and CSPP-L in cells using a panel of GFP-tagged UBR5 variants (wild-type and functional domain mutants ΔUBR, ΔMLLE and ΔHECT - as detailed in Fig 3A). Co-immunoprecipitation using GFP-UBR5 in HEK293T cells detected an interaction between UBR5 and endogenous CSPP-L, while no interaction was observed using GFP-only control (Fig. 3B, S3B). Mutation of the known functional domains in UBR5 (UBR-box (ΔUBR – UBR5^W1235L^), MLLE domain (ΔMLLE – UBR5^Y2402A/K2415A^) or HECT active site (ΔHECT – UBR5^C2768A^) had no detectable effect on the immunoprecipitation of CSPP-L (Fig 3B), suggesting that the interaction between these proteins is independent of these domains in UBR5 (transfection efficiency for these experiments is detailed in Fig S3A). The interaction between UBR5 and CSPP-L was confirmed using GFP-labelled CSPP-L to immunoprecipitate endogenous UBR5 from HEK293T cells, but not with GFP-only control (Fig 3C). A significant amount of CSPP-L was still detectable in the non-bound fraction of GFP-UBR5 pull-downs (Fig. 3B), suggesting that UBR5 is not sequestering the complete pool of cellular CSPP-L in this assay.

**Figure 3:**
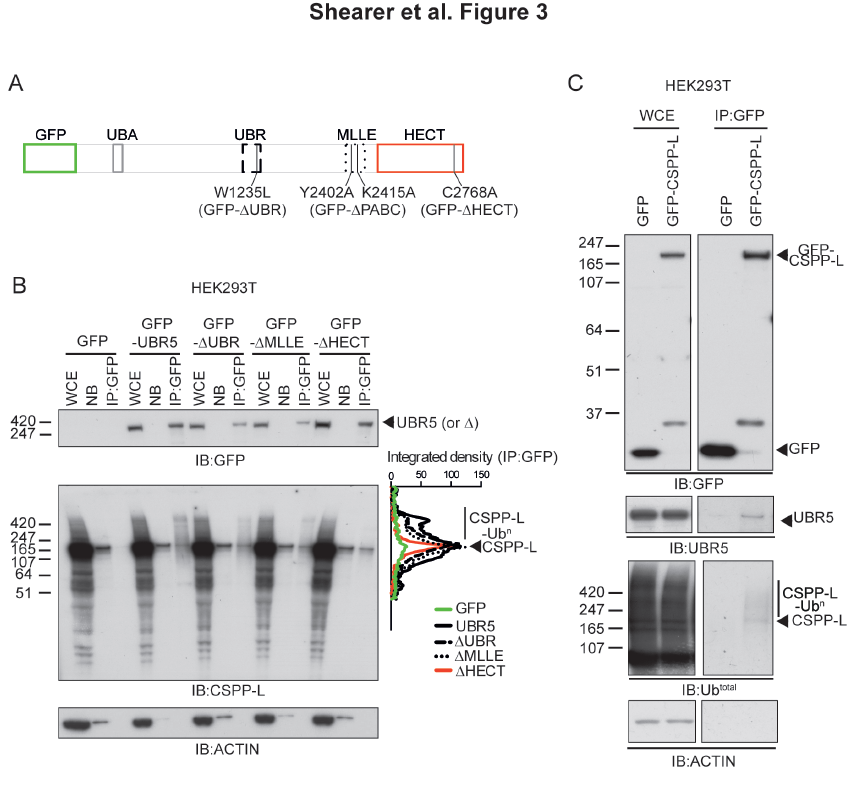
UBR5 binds and poly-ubiquitylates CSPP-L. **A**. Schematic showing structure of UBR5 protein and position of known functional domains and amino acid changes of A mutants. **B**. Co-immunoprecipitation and immunoblot analysis of GFP-UBR5 (and functional mutants) transfected into HEK293T cells. GFP-UBR5 co-immunoprecipitates endogenous ubiquitylated CSPP-L. Poly-ubiquitylation of CSPP-L is quantified by integrated density plot showing relative size shift. Note the loss of poly-ubiquitylated CSPP-L binding ΔHECT mutant. GFP-UBR5 and A mutants are predicted to be approximately 338kDa. Transfections performed 24 hours prior to harvest. WCE - Whole cell extract. NB - Non-bound fraction. (4% of input). IP - Immunoprecipitation. **C**. Co-immunoprecipitation and immunoblot analysis of GFP-CSPP-L transfected into HEK293T cells. GFP-CSPP-L binds endogenous UBR5. Endogenous Ub is detected covalently bound to GFP-CSPP-L. GFP-CSPP-L is predicted to be approximately 170kDa.

The presence of higher molecular weight smears on CSPP-L western blots following pull-down with GFP-UBR5 (Fig 3B) suggested the presence of poly-ubiquitylated forms of CSPP-L in HEK-293T cells. Probing with an anti-Ub antibody following pulldown of GFP-CSPP-L confirmed the presence of ubiquitylated CSPP-L (CSPP-L-Ub) in HEK-293T cells (Fig 3C). These presumptive poly-ubiquitylated forms of CSPP-L were attenuated in pull-downs using the ligase-dead form of UBR5 (ΔHECT), suggesting direct ubiquitylation of CSPP-L by UBR5 (Fig 3B, S3B). Importantly, this experiment also confirmed the presence of ubiquitylated forms of endogenous CSPP-L (i.e. not GFP labelled). Over-expression of UBR5 did not alter total cellular CSPP-L levels (Fig S3B), suggesting that UBR5-mediated ubiquitylation of CSPP-L is not targeting the protein for degradation by the proteasome.

### UBR5 interacts with CSPP-L at the centrosome

We sought to further characterise the UBR5 and CSPP-L interaction in a cellular context using protein complementation assays to define subcellular localisation. Bi-molecular fluorescence complementation (BiFC) analysis (Fig. 4A) indicated the presence of protein-protein interactions between CSPP-L and UBR5 in discrete perinuclear foci in HEK293T cells, consistent with the primary sub-cellular localisation of CSPP-L in centrosomes (white arrows, Fig 4A). A similar subcellular distribution was observed for BiFC signal generated by an interaction between CSPP-L and Ub (Fig. 4A), suggesting that CSPP-L is ubiquitylated at the centrosome and consistent with our results above (Fig 3). A positive control assay between UBR5 and Ub showed strong nuclear and cytoplasmic foci (Fig. 4A) consistent with the previously described localisation of UBR5 to the nucleus and cytoplasm (Fig S3A) (Fuja et al., 2004). Correct expression of V1 or V2 tagged BiFC fusion proteins was confirmed by immunoblot for both the expression protein and respective Venus fluorescent protein fragment (Fig 4B).

**Figure 4:**
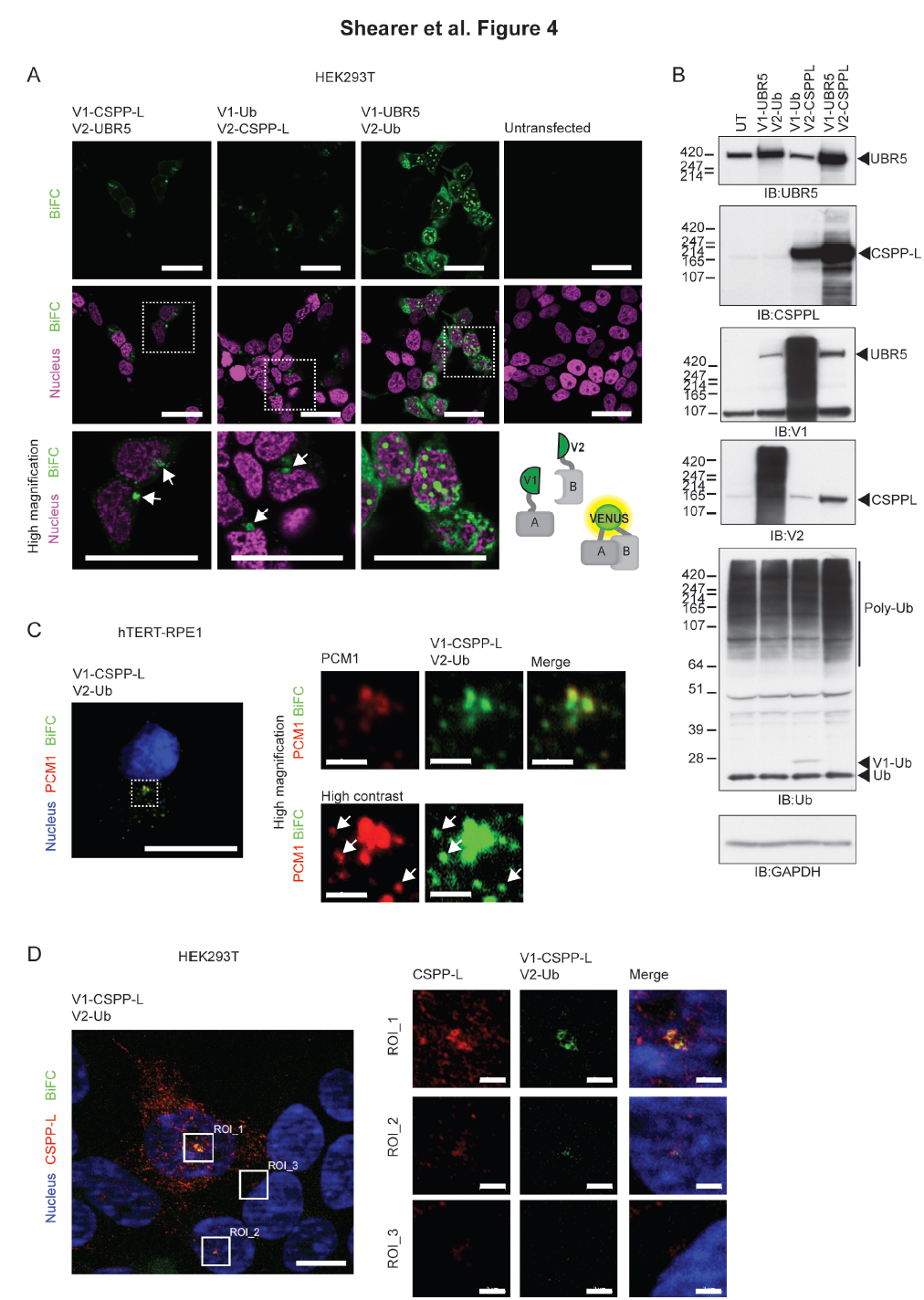
UBR5 interacts with CSPP-L at the centrosome. **A**. Left to right, CSPP-L interacts with UBR5 in large foci adjacent to the nucleus (arrows). CSPP-L interacts with Ub in large foci adjacent to the nucleus (arrows). Interaction between UBR5 and Ub in HEK293T cells shows nuclear and cytoplasmic localization, with large foci of UBR5 and Ub in the nucleus. Untransfected cells show no visible signal. Nuclear marker is H2B-mCherry (magenta). High powered inset fields indicated by dashed box (bar = 30μm) and schematic of BiFC analysis for protein-protein interactions included. **B**. Immunoblot data showing correct production of fusion proteins in BiFC assay. Expected fusion protein sizes are: V2-CSPP-L 152kDa, V1-UBR5 329kDa, V1-Ub 28kDa, V2-Ub 19kDa. **C**. CSPP-L/Ub interacting pairs co-localise with PCM1 (marker of centrosome and centriolar satellites) in hTERT-RPE1 cells. Bar = 20μm. High magnification bar = 2μm. **D**. High level expression of CSPP-L/Ub BiFC vectors shows CSPP-L and Ub interaction is confined to the centrosomal region, despite strong detection of CSPP-L at the microtubules in HEK293T cells. Region of interest (ROI) 1 shows relatively high BiFC pair expression, ROI 2 shows relatively low BiFC pair expression and ROI 3 shows no BiFC pair expression. Bar = 10μm. High magnification of ROI bar = 2μm.

We next used BiFC to examine ubiquitylation of CSPP-L interaction pairs in hTERT-RPE1 cells and co-stained for PCM1 (which marks the centrosome and satellites) to confirm detailed localisation. BiFC signal indicating Ub:CSPP-L was detected at the centrosome (Fig 4C) and surrounding satellites structures (Fig 4C, white arrows), indicating that the perinuclear BiFC signal observed in HEK293T cells (Fig 4A) was indeed localised at the centrosome and surrounding satellite material. These data indicate that CSPP-L is ubiquitylated at the centrosome and surrounding satellites.

### UBR5 regulates recruitment of CSPP-L to the centrosome and centriolar satellites

Consistent with effects of UBR5 overexpression (Fig. 3B), we observed no change in total cellular CSPP-L following UBR5 depletion by shRNA (Fig 5A). Further, we did not observe accumulation of ubiquitylated CSPP-L following treatment with the proteasome inhibitor MG-132 (Fig. S3B). Together, these data strongly indicate that UBR5-mediated ubiquitylation of CSPP-L is not targeting the protein for degradation. Analysis of GFP-CSPP-L pulldowns using chain-specific Ub antibodies indicated the presence of both K48 and K63 linked polyubiquitin chains on CSPP-L (Fig 5B). Nondegrading K63 linked poly-ubiquitin chains have been implicated in endocytosis trafficking and signal transduction (Ikeda and Dikic, 2008).

**Figure 5:**
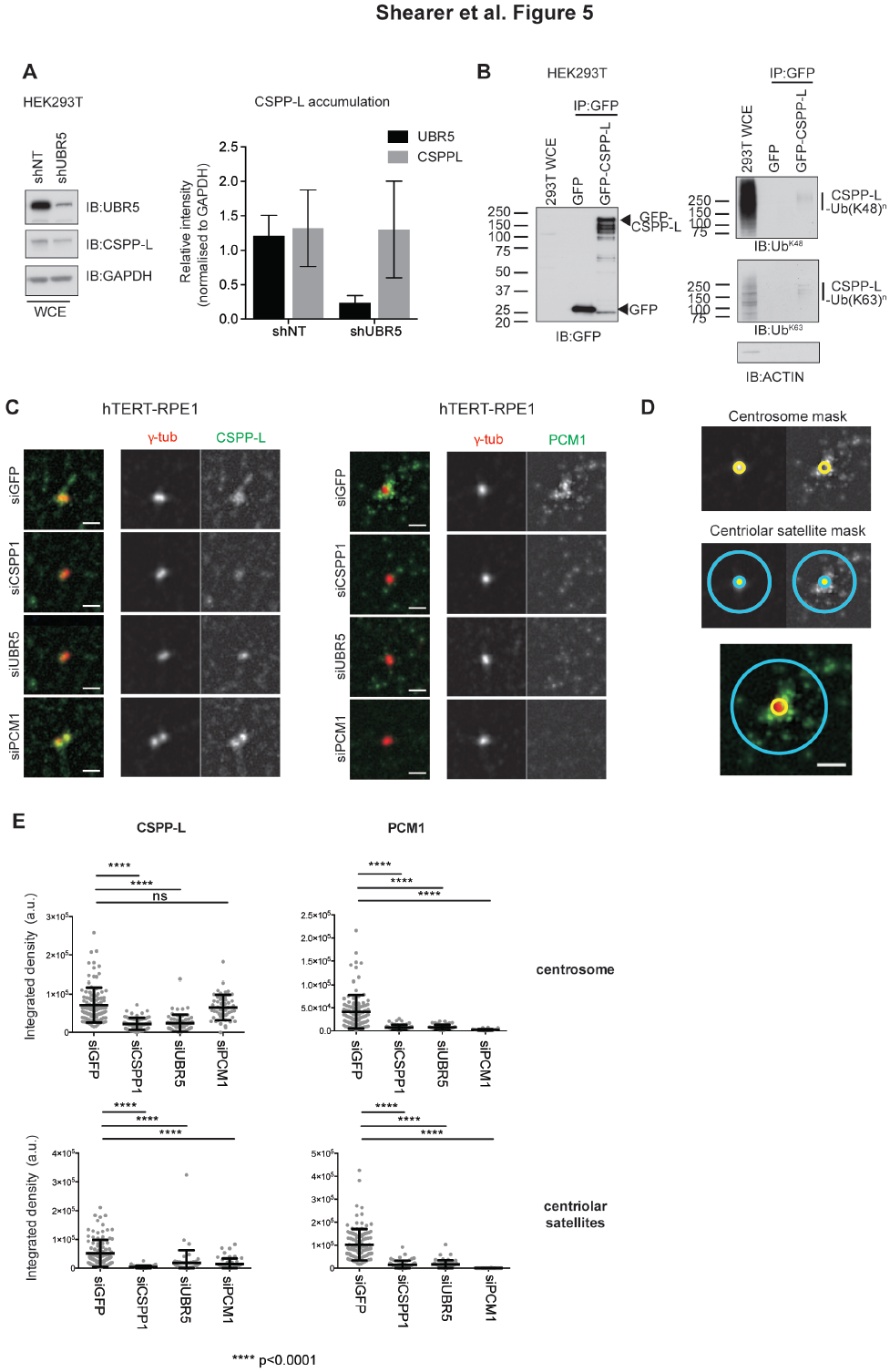
UBR5 maintains CSPP-L at the centrosome/centriolar satellites, and is required for centriolar satellite stability. **A**. Depletion of UBR5 by shRNA in HEK293T cells does not decrease CSPP-L levels. shRNA induced with 1μg/ml DOX 48 hours prior to harvest. Data summary representative result from 4 independent experiments, n=6 replicates per experiment. **B**. Immuniprecipitation of GFP-CSPP-L co-immunoprecipitates CSPP-L-bound lysine-48 (K48) linked Ub chains and lysine-63 (K63) linked Ub chains. Transfections performed 24 hours prior to harvest. **C**. Depletion of UBR5 (but not PCM1) causes dispersion of centrosomal CSPP-L in RPE1 cells. **D**. Centrosome/centriolar satellite mask **E**. Quantitation of loss of CSPP-L/PCM1 at the centrosome according to the mask described in (**D**) At least 30 cells were scored per sample and statistical analysis performed using two-tailed t-test.

Interestingly, transiently co-expressed V1-CSPP-L and V2-Ub fusion proteins resulted in BiFC signal that co-localized with PCM1 in hTERT-RPE1 cells (Fig 4C), indicating presence of ubiquitylated CSPP-L at centriolar satellites. Moreover, the V1-CSPP-L:V2-Ub BiFC signal was confined to centriolar satellites alone, since excessive ectopic (and endogenous) CSPP-L was detected at MTs by a CSPP-L specific antibody (Fig4D).

Depletion of UBR5 diminished localisation of CSPP-L to the centrosome (Fig. 5C, E), and adjacent satellites as determined by a pericentriolar quantitation mask (Fig 5D). This effect was independent of disruption of centriolar satellites, as depletion of PCM1 did not prevent aggregation of CSPP-L at the centrosome (Fig. 5C (and Sternelalm et al., under review)). Loss of PCM1 is known to disrupt centriolar satellite stability and ciliogenesis (Wang et al., 2016), and accordingly PCM1 disruption depletes centriolar satellite accumulation of CSPP-L (Fig 5C). It is apparent that CSPP-L and UBR5 are integral to satellite stability, as depletion of either almost completely attenuates detectable PCM1 directly adjacent to the centrosome (Fig 5C).

## Discussion

We have demonstrated a novel function for the E3 Ub ligase UBR5 in regulation of ciliogenesis via maintenance of centriolar satellite stability. UBR5 mediates this effect via a novel protein-protein interaction between UBR5 and the CSPP-L isoform of CSPP1, predominantly at the centrosome and surrounding centriolar satellites. CSPP-L is an established positive regulator of ciliogenesis (Patzke et al., 2010) and a prominent cilia-localising protein (Mick et al., 2015). Cilium assembly is tightly linked to exit from mitosis into G1 phase (Rieder et al., 1979). CSPP-L is increasingly recruited to the centrosome during G2/Prophase in cell cycle progression, and detected on spindle MTs during mitosis, concomitant with centriolar satellite dispersal (Patzke et al., 2006, Kim et al., 2016). This cell cycle dependent localization of CSPP-L to the primary cilium and spindle apparatus is likely regulated via post-translational regulatory mechanisms. Indeed, the UPS is a known regulator of centriolar satellite stability (Shearer and Saunders, 2016) and we observed that UBR5-mediated ubiquitylation of CSPP-L is necessary to stabilise not only centriolar satellite organization, but also CSPP-L’s centrosomal localization. Hence, UBR5 is important for the maintenance of centriolar satellite stability and centrosome integrity.

Another E3 ubiquitin ligase, MIB1, is also known to regulate ciliogenesis via ubiquitin-mediated regulation of PCM1 function (Villumsen et al., 2013, Wang et al., 2016). We observed a weak interaction between UBR5 and MIB1, but not between UBR5 and PCM1. This may reflect a general accumulation of UBR5 around the centrosome, as indicated by the interaction of UBR5 with γ-Tubulin, or may also indicate large multi-protein complexes arising at centriolar satellites. However, as these would also accumulate PCM1 - a major component of centriolar satellites (Kubo et al., 1999) - this explanation for the observed weak UBR5-MIB1 interaction is less probable.

Even though we detected ubiquitylation of CSPP-L in association with UBR5, siRNA-mediated depletion of UBR5 did not alter cellular levels of CSPP-L, suggesting that UBR5 is not targeting CSPP-L for proteasomal degradation. BiFC experiments indicate that ubiquitinylated CSPP-L is primarily confined to centriolar satellites, while excessive and non-ubiquitylated CSPP-L (i.e. not detected by BiFC for Ub:CSPP-L) localizes to MTs (Fig 4D). Notably, both K48 and K63 linked poly-Ub conjugated forms of CSPP-L were detected interacting with UBR5, indicating the presence of non-degrading Ub signalling events. UBR5 is known to assemble non-degrading ubiquitin chains on β-catenin (Hay-Koren et al., 2011) and ATMIN (Zhang et al., 2014). Non-degrading Ub signalling is also involved in other aspects of centriolar satellite maintenance. For example, MIB1 monoubiquitylates PCM1, AZI and CEP290 in the absence of UV cellular stress and maintain these proteins in an inactive form until monoubiquitylation is reversed (Villumsen et al., 2013). However, another study found PCM1 to be degraded by MIB1 mediated polyubiquitylation (Wang et al., 2016) with discrepancies in results likely due to different methodologies used in each study (Shearer and Saunders, 2016). Depletion of either CSPP-L or UBR5 disrupted centriolar satellite organization (as defined by PCM1), indicating that UBR5 mediated ubiquitylation of CSPP-L promotes localization of CSPP-L to centriolar satellites. The UBR5/CSPP-L interplay is thus one regulatory mechanism controlling the timely release and activity of ciliogenesis promoting factors, including CSPP-L itself.

Primary cilia are an important component of the Hh signal transduction pathway during development (Rohatgi et al., 2007, Goetz and Anderson, 2010) and autocrine Hh signalling is reactivated in some cancer types (Kubo et al., 2004, Liu et al., 2014, Ertao et al., 2016). Several studies have implicated UBR5 as a modulator of Hh signalling and a variety of model organisms with UBR5 mutations display developmental defect phenotypes. Hence, it is worth considering a potential underlying role for UBR5 in Hh signalling via the novel role in regulating ciliogenesis. Mutations in the *Drosophila* ortholog of UBR5 (Hyd) display a range of severe developmental phenotypes (reviewed in (Shearer and Saunders, 2016)). Specifically, Hyd directly regulates Hh and Dpp signalling (Lee et al., 2002), and can also indirectly affect Hh signal transduction by regulating Ci promoter binding (Wang et al., 2014). UBR5 null mouse embryos die at mid gestation due to failure of yolk sac vascular development (Saunders et al., 2004). Coincidently, this timing corresponds to the developmental stage at which primary cilia firstly appear on epiblast-derived mesothelial and endothelial cells (Bangs et al., 2015). Conditional deletion of *Ubr5* in the early embryonic limb-bud mesenchyme of mice resulted in decreased Hh ligand production and decreased Hh pathway activity (Kinsella et al., 2016). It was not determined whether these effects on Hh signaling were direct (cell autonomous) or indirect (non-cell-autonomous) and so the underlying mechanism for this effect remains elusive. UBR5 has not been shown to directly bind Hh pathway components in human cells apart from GLI2 (Moncrieff et al., 2015), however multiple Hh pathway components including PTCH1, GLI1 and HHIP are direct transcriptional targets of active Hh signalling (Gupta et al., 2010). It cannot be excluded that a reduction in general Hh activity with reduced UBR5 may indicate a more general disruption in signal transduction caused by failed ciliogenesis.

From a disease perspective, UBR5 has not been specifically examined in the context of ciliopathies. Rare *UBR5* missense mutations have been linked to familial epilepsy (Kato et al., 2012) and cilia have been implicated in some forms of epilepsy (Delgado-Escueta, 2007) but this link is speculative. Exome sequencing data from the Exome Aggregation Consortium (ExAC) demonstrates a very low LoF mutation rate for UBR5 in healthy somatic tissue. Only 4 LoF UBR5 variants were observed, with a very low frequency (Lek et al., 2016). These indicate strong selective pressure against deactivating UBR5 mutations and support a critical role for UBR5 in human development, consistent with observations in model organisms.

Correlated expression of *CSPP1* and *UBR5* mRNA in human cancer cell lines and primary breast cancer gives additional relevance of CSPP-L and UBR5 in normal and transformed mammary epithelium. However, further work is required to understand the putative impact of epithelial lineage-specific expression of CSPP1 isoforms (which can’t be distinguished by mRNA expression data) and their co-expression and interaction with UBR5 (Sternemalm et al., 2014). However, it is interesting to note that ciliary Hh signaling controls branching morphology of mammary gland xenografts (McDermott et al., 2010) and the Hh signalling pathway is a growth-promoting factor for breast cancer subgroups (Kubo et al., 2004, O’Toole et al., 2011). Future analysis may focus on determining putative correlations between UBR5/CSPP-L controlled cilia formation and mammary epithelial cell differentiation and transformation.

In summary, we have demonstrated a highly novel role for the E3 Ub ligase UBR5 in primary cilia maintenance/formation through ubiquitylation of CSPP-L, with potential implications for understanding the molecular basis of key signalling pathways in development and disease.

## Acknowledgements

This research was supported by funding from the National Health and Medical Research Council of Australia (project grant GNT1052963), Cancer Institute NSW 10/FRL/3-02], Mostyn Family Foundation, and the Australian Government Department of Innovation, Industry, Science and Research (DIISR). RFS is the recipient of the Baxter Family Scholarship. SP is the recipient of a Career Development Fellowship from the Norwegian Cancer Society. DNS and AB are recipients of the Patricia Helen Guest Fellowship.

